# Radiances of Cerenkov-Emitting Isotopes on the IVIS

**DOI:** 10.1101/2023.01.18.524625

**Authors:** Edwin C. Pratt, Travis M. Shaffer, David Bauer, Jason S. Lewis, Jan Grimm

**Affiliations:** Department of Pharmacology, Weill Cornell Graduate School; Department of Radiology, Memorial Sloan Kettering Cancer Center; Department of Molecular Pharmacology, Memorial Sloan Kettering Cancer Center

## Abstract

Cerenkov (or Cherenkov) luminescence occurs when charged particles exceed the phase velocity of a given medium. Cerenkov has gained interest in preclinical space as well as in clinical trials for optical visualization of numerous radionuclides. However, Cerenkov intensity has to be inferred from alternative databases with energy emission spectra, or theoretical fluence estimates. Here we present the largest experimental dataset of Cerenkov emitting isotopes recorded using the IVIS optical imaging system. We report Cerenkov measurements spanning orders of magnitude normalized to the activity concentration for 21 Cerenkov emitting isotopes, covering electron, alpha, beta minus, and positron emissions. Isotopes measured include Carbon-11, Fluorine-18, Phosphorous-32, Scandium-47, Copper-64, Copper-67, Gallium-68, Arsenic-72, Bromine-76, Yttrium-86, Zirconium-89, Yttrium-90, Iodine-124, Iodine-131, Cerium-134, Lutetium-177, Lead-203, Lead-212, Radium-223, Actinium-225, and Thorium-227. We hope this updating resource will serve as a rank ordering for comparing isotopes for Cerenkov luminescence in the visible window and serve as a rule of thumb for comparing Cerenkov intensities in vitro and in vivo.

**Methods:** All Cerenkov emitting radionuclides were either produced at Memorial Sloan Kettering Cancer Center (Carbon-11, ^11^C; Fluorine-18, ^18^F; Iodine-124, ^124^I), from commercial sources such as Perkin Elmer (Phosphorous-32, ^32^P; Yttrium-90, ^90^Y), Bayer (Radium-223, ^223^Ra, Xofigo), 3D-Imaging (Zirconium-89, ^89^Zr), Nuclear Diagnostic Products (Iodine-131, ^131^I), or from academic collaborators at Washington University at St. Louis (Copper-64, ^64^Cu), University of Wisconsin (Bromine-76, ^76^Br), MD Anderson Cancer Center (Yttrium-86, ^86^Y), Brookhaven National Laboratory (Arsenic-72, ^72^As; Thorium-227, ^227^Th), or Oak Ridge National Laboratory (Cerium-134, ^134^Ce, Actinium-225, ^225^Ac), and Viewpoint Molecular Targeting (Lead-203, ^203^Pb; Lead 212, ^212^Pb). All isotopes were diluted in triplicate on a black bottomed corning 96 well plate to several activity concentrations ranging from 0.1-250 μCi in 100-200 μL of Phosphate Buffered Saline. Cerenkov imaging was acquired on a single Perkin-Elmer Spectrum In-Vivo Imaging System (IVIS) at field of view c with exposures ranging up to 15 minutes or lower provided no part of the image intensity was saturated, or that the activity significantly changed during the exposure. Experimental radiances on the IVIS were calculated from regions of interest drown over each 96 well, and then normalized for the activity present in the well, and the volume the isotope was diluted into.

## Introduction and Results

Cerenkov Luminescence occurs when a charged particle exceeds the speed of light in a given medium^1^. Cerenkov Luminescence Imaging was first used in a biomedical research setting back in 2009 with the availability of the IVIS system^2^. Subsequent work has utilized Cerenkov luminescence by dye^3^ or nanoparticle mediated^4,5^ energy transfer, light base therapy^6^, in addition to direct conjugates to numerous molecular imaging agents^7^. Previous studies have shown fluence estimates in tissue^8^ as well as experimental results comparing select sets of isotopes^4,9^ with many applications reviewed previously^10,11^ with recent updates from a clinical trial^12^. Through the course of numerous radiochemistry experiments, the following twenty-two Cerenkov radiances have been acquired and summarized below in **Figure 1a** with isotopes rank ordered by mean measured radiance that is normalized to the activity concentration. Furthermore Cerenkov radiances are provided in **Figure 1 b-d** by emission type, covering electron, alpha, beta minus, and positron emission for clearer comparison within emission groups. Within the alpha emitting panel of **Figure 1b**, alpha emissions are further speciated as haven been acquired after recent purification, or after sufficient time to allow equilibrium with the daughter radionuclides. Emissions presented here agree with rank ordering of beta emission profiles found in the Nuclear Data Sheets.

**Figure 1.**
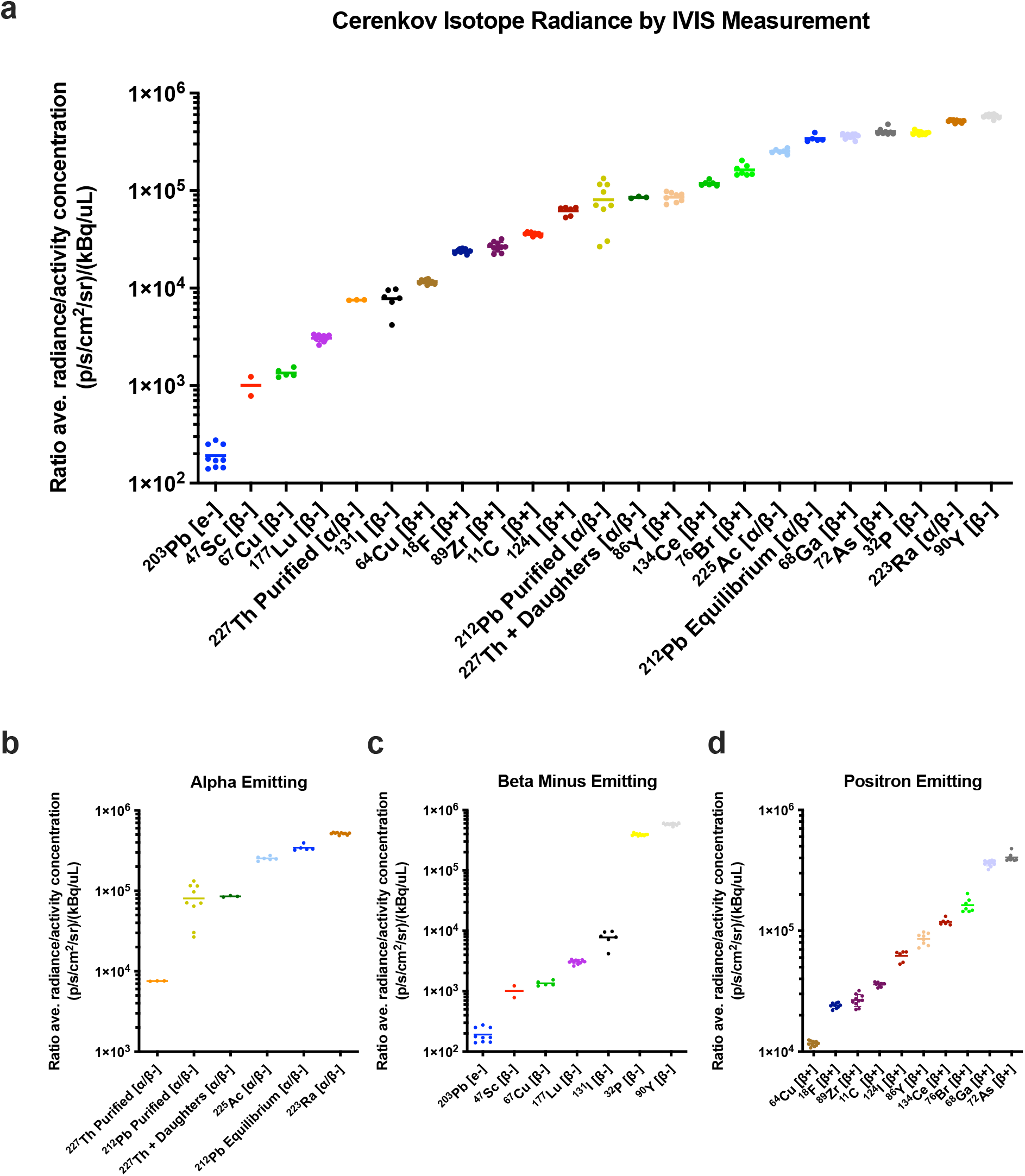
Experimental Cerenkov radiances acquired on the IVIS Spectrum for numerous clinical radionuclides. **a**, Rank ordering of Cerenkov isotopes measured on the IVIS show a broad range of radiances between 1E2 and 1E6 p/s/cm_2_/sr. **b**, Alpha emitting radionuclides rank ordered and measured either in equilibrium, or right after separation from decay daughters. **c**, beta minus emitting or **d**, positron emitting groupings to compare isotopes within each emission type.

## Discussion

At the highest end, ^90^Y and ^223^Ra, ^32^P and ^72^As represent the strongest Cerenkov emitting isotopes per unit of activity per volume. On the other end, isotopes such as ^67^Cu, ^47^Sc, and ^177^Lu represent therapeutic isotopes with the lowest Cerenkov radiance due to their lower beta energy spectrum. The electron emitting ^203^Pb represents the lowest intensity Cerenkov emission, with only ∼5% of fast electrons emitted sufficient for Cerenkov luminescence in water. In the middle range, many of the more commonly used positron emitting isotopes such as ^11^C, ^18^F, ^64^Cu, and ^89^Zr reside between 1E4 and 3E4 on the graph, showing similar efficiency per unit of activity compared to the lowest or highest emitting radionuclides on the order of 1E2 and 1E6, respectively.

## Conclusion

This work presents the culmination of numerous Cerenkov experiments, each measuring only the isotope present on the same IVIS system. Here over 20 isotopes, with and without decay chain daughters can be compared side by side for their experimental Cerenkov luminescence efficiency. Cerenkov isotope emissions range between 1E2 and 1E6 p/s/cm^2^/sr per kBq/μL on the IVIS Spectrum with an open filter, while the most common clinically used radioisotopes such as ^11^C, ^18^F, ^64^Cu, and ^89^Zr emit near 1E4 p/s/cm^2^/sr per kBq/μL. Together this work serves as a guide to compare Cerenkov intensities between isotopes available for molecular imaging and theranostic use.

## Acknowledgements

We would like to acknowledge many of the fellow Grimm and Lewis, and Zeglis Lab members who through various scientific needs, brought all of the isotopes to Memorial Sloan Kettering Cancer Center and graciously let us measure fractions of the precious isotope supply for this Cerenkov luminescence study. We would also like to thank Pat Zanzonico and Valerie Longo of the MSKCC Small animal core for the support and use of the IVIS spectrum instrument for Cerenkov luminescence imaging. R01 CA215700 (to J.G.), S10 OD016207-01 (to Pat Zanzonico, MSKCC), and P30 CA08748 (to Craig Thompson MSKCC). E.C.P. is currently supported by the National Institutes of Health F32 CA268912-01.

## Notes

### Competing Interest Statement

The authors have declared no competing interest.

